# Regulation of apico-basolateral trafficking polarity of homologous Copper-ATPases ATP7A and ATP7B

**DOI:** 10.1101/2023.04.19.537613

**Authors:** Ruturaj, Monalisa Mishra, Soumyendu Saha, Saptarshi Maji, Enrique Rodriguez-Boulan, Ryan Schreiner, Arnab Gupta

## Abstract

We suggest a model of apico-basolateral sorting in polarized epithelia using homologous Cu-ATPases as membrane cargoes. In polarized epithelia, upon copper treatment, homologous copper-ATPases ATP7A and ATP7B traffic from trans-Golgi network (TGN) to basolateral and apical membranes respectively. We characterized sorting pathways of Cu-ATPases between TGN and plasma-membrane and identified the machinery involved. ATP7A and ATP7B reside on distinct domains of TGN and in high copper, ATP7A traffics directly to basolateral membrane, whereas ATP7B traverses common-recycling, apical-sorting and apical-recycling endosomes *en-route* to apical membrane. Mass-spectrometry identified regulatory partners of ATP7A and ATP7B that include Adaptor Protein-1 complex. Upon knocking-out pan-AP-1, sorting of both copper-ATPases are disrupted. ATP7A loses polarity and localizes on both apical and basolateral surfaces in high copper. Contrastingly, ATP7B loses TGN-retention but retains apical polarity that becomes copper-independent. Using isoform-specific knockouts, we found that AP-1A provides directionality and TGN-retention for both Cu-ATPases, whereas, AP-1B governs polarized trafficking of ATP7B solely. Trafficking phenotypes of Wilson disease-causing ATP7B mutants that disrupts putative ATP7B-AP1 interaction further substantiates the role of AP-1 in apical sorting of ATP7B.

**Summary statement:** The Adapter Protein-1 (AP-1) isoforms AP-1A and AP-1B governs the apico-basolateral trafficking polarities of the homologous Copper-ATPases, ATP7A and ATP7B at the trans-Golgi network.

## Introduction

Copper is a crucial micro-nutrient for all eukaryotic organisms ^1, 2^. Copper is a redox-active metal ^3, 4^. Its intracellular levels need to be tightly regulated as copper accumulation causes severe toxicity ^5^. Cellular copper is primarily regulated by Golgi localizing copper transporting ATPases, ATP7A and ATP7B, highly homologous P-type ATPases that pump copper into the lumen of the TGN, where various proteins utilize it as co-factor for their maturation ^6–8^. Upon elevated copper, these Cu-ATPases are redistributed by vesicular traffic from TGN to the plasma membrane, where they export excess copper out of the cell ^9–11^. Mutations in ATP7A, ubiquitously expressed and abundant in enterocytes, lead to systemic deficiency of copper causing Menkes disease ^12^. In contrast, mutations in ATP7B lead to copper accumulation in liver, kidney and brain leading to Wilson disease, characterized by liver cirrhosis and neurological symptoms ^13, 14^.

Lower organisms that precede chordate evolution have only one copper ATPase. Evolutionary divergence from a single copper ATPase to ATP7A and ATP7B is first observed in fish ^15^. Human ATP7A and ATP7B share ∼65% sequence homology, with highly conserved domains that include the cytoplasmic amino terminus harbouring six copper-binding domains containing the sequence MXCXXC, the nucleotide-binding domain, the phosphorylation domain, the actuator domains and the carboxy terminus ^16^. Under normal physiological conditions, as cellular copper increases, the six metal-binding domains in the NH_2_-terminal sequester copper. This causes structural changes in the proteins that promote their redistribution to the plasma membrane to export excess intracellular copper ^17, 18^. In polarized epithelial cells exposed to elevated copper, both transporters leave the TGN with ATP7A directed to the basolateral surface and ATP7B to the apical surface ^19, 20^. Studies carried out on the epithelial cell line MDCK and other epithelial cell lines have identified the TGN as the compartment where apical and basolateral plasma membrane proteins are segregated into distinct routes to the plasma membrane ^21–27^. During transport to the cell surface, some plasma membrane proteins may traffic through endosomal compartments, such as common recycling endosomes (CRE), apical recycling endosomes (ARE) and apical sorting endosomes (ASE), where additional sorting events may take place ^28–32^. Transport along these routes is directed by apical signals such as N-glycans ^33^, O-glycans ^34^; GPI anchors ^35, 36^; and basolateral signals that resemble in structure with tyrosine and di-leucine endocytic motifs ^37^. A variety of molecules have been postulated to mediate these trafficking processes, such as Clathrin ^38^, Clathrin adaptors AP-1A and AP-1B ^30, 39–43^, microtubule motors ^29, 44, 45^ and regulators of the actin cytoskeleton ^46–48^.

Limited trafficking studies have been carried out on the polarized trafficking of ATP7A and ATP7B. ATP7A has been mostly studied in non-epithelial cell models like HeLa and HEK ^49–51^. Hubbard and coworkers have utilized the liver epithelial cell line WIF-B as a model to study apical trafficking signals in ATP7B. Guo et al found that the Wilson disease-causing mutation ATP7B-N41S results in the mislocalization of ATP7B to the basolateral surface ^47^. The N41S mutation lies within a stretch of 9 amino acids that was determined to be essential for apical targeting of ATP7B in polarized hepatocytes; the deletion of which resulted in basolateral targeting of ATP7B ^47^. In neurons, the YXXФ motif at the C-terminus of ATP7B was shown to be critical in maintaining somatodendritic polarity and TGN localization ^52^. The *bona fide* clathrin-interacting dileucine motif within the DKSWLLL stretch on the C-terminus of ATP7B has also been shown to be crucial for its copper-dependent trafficking ^53^. Similarly, a di-leucine motif at the C-terminus of ATP7A was found to be important for its internalization and retrograde transport from the plasma membrane to TGN ^51^. Myosin Vb and its effector Rab-GTPase Rab11 were shown to be essential for the apical trafficking of ATP7B ^48^. Fundamental questions remain unanswered on the differential regulation, directionality and the sorting compartments involved in the polarized trafficking of ATP7A and ATP7B.

We investigated the intracellular pathways, signals and mechanisms directing ATP7A and ATP7B to opposite poles of epithelial cells using the Madin Darby Canine Kidney (MDCK) cell line. These two proteins are ideally suited for these studies as they are highly homologous and exhibit synchronized trafficking in response to the same ligand i.e., copper. The absence or chelation of copper naturally acts as their Golgi-exit blocker as they are primarily Golgi-resident proteins. Our results identify intracellular compartments involved in the trafficking of these two proteins and demonstrate the role of Clathrin adaptors AP-1A and AP-1B in their polarized sorting.

## Results

### ATP7A and ATP7B traffic to basolateral and apical surfaces respectively in polarized MDCK cells

ATP7A and ATP7B are highly homologous proteins with conserved functional domains. However, they exhibit sequence dissimilarity in two specific regions, i.e., at the extreme amino-terminus and a flexible unstructured loop at the nucleotide-binding domain (Fig. 1A). Some cell types co-express both the copper-ATPases. Both Menkes disease, as well as Wilson disease patients, exhibit copper accumulation in tubules of the nephron ^54^. Barnes and co-workers have shown that HEK293, a cell line derived from human embryonic kidney expresses both ATP7A and ATP7B ^55^. We found that MDCK cells express transcripts for both the copper transporters (Fig. 1B). We have previously shown that Cu levels do not affect the endogenous expression levels of ATP7A and ATP7B, which corroborates findings from other groups (Das et al., 2022; Barnes et al., 2009; Leonhardt et al., 2009). Due to the unavailability of commercial antibodies against dog ATP7A and ATP7B, we expressed fluorescently tagged ATP7A and ATP7B (i.e., mKO2-HA-ATP7A and eGFP-ATP7B). mKO2 is a monomeric orange fluorescent protein with an excitation peak of 552nm and an emission peak of 565nm. We used salt of Tetrathiomolybdate (TTM: 25µM) or Bathocuproinedisulfonic acid (BCS: 50µM) as copper chelators to mimic low copper conditions; CuCl_2_ (10-50µM) treatment in culture media was used to elevate intracellular copper levels. Using ICP-MS copper levels in basal media was determined to be ∼0.7±0.2µM.

**Figure 1.**
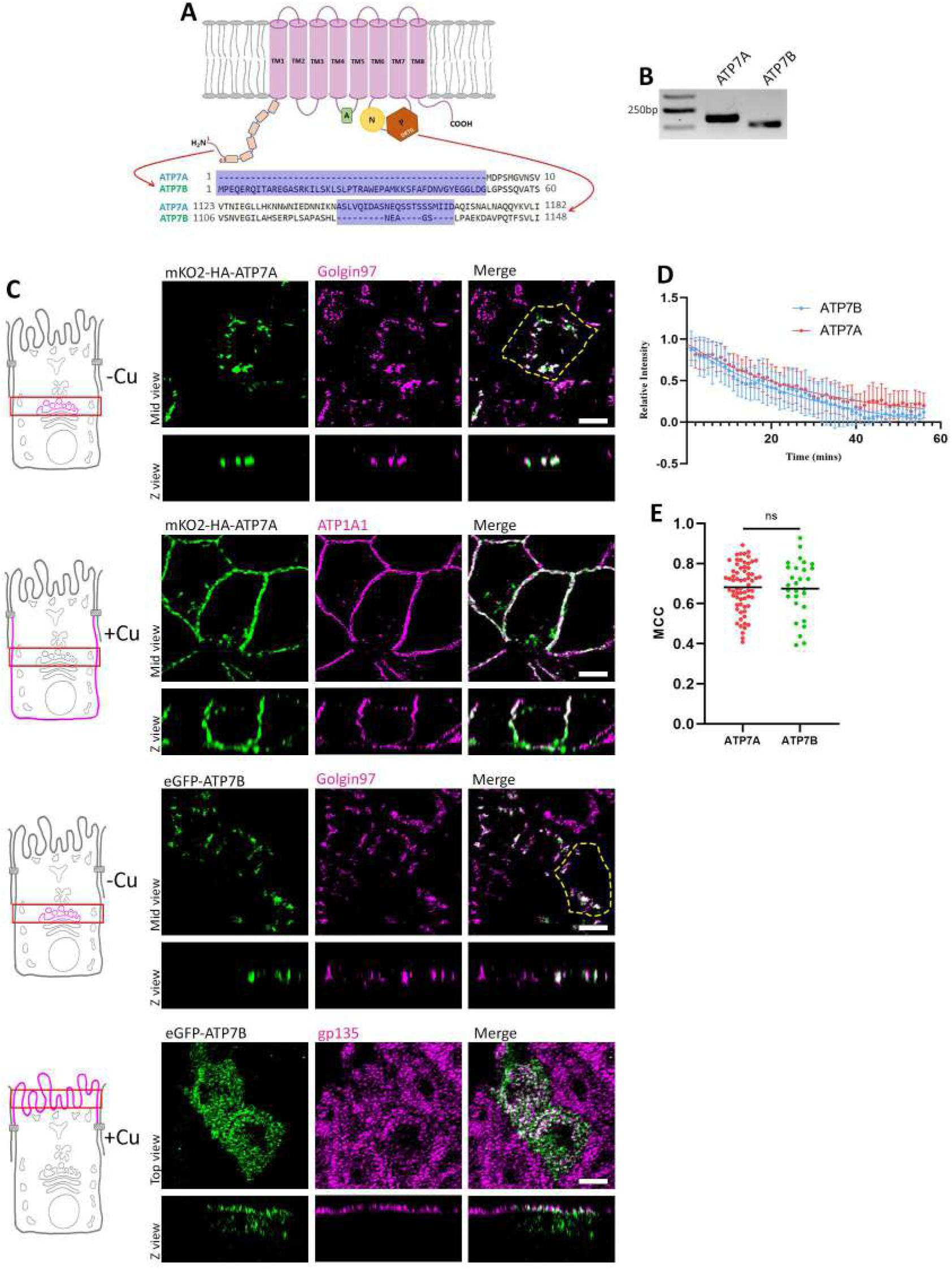
ATP7A and ATP7B are homologous and are expressed in MDCK cells but traffic to different surfaces in response to copper. **(A)** Illustration showing the structure of Copper ATPases with major sequence differences between ATPA and ATP7B drawn out. **(B)** Transcripts of dog ATP7A and ATP7B in polarized MDCK cells, shows both the transporters are expressed in MDCK. **(C)** Polarized MDCK cells showing localization of transfected mKO2-HA-ATP7A and eGFP-ATP7B in copper-depleted and copper-treated conditions. Under copper-depleted conditions, both proteins localize at the TGN marked by Golgin97. Upon copper treatment, ATP7A traffic to the basolateral surface (marked by ATP1A1) and ATP7B traffics to the apical surface (marked by gp135). **(D)** Fitted curve of decreased pixel count of both the Cu ATPases, mKO2-HA-ATP7A and eGFP-ATP7B, in response to copper, shows similar dispersion as their exit rate (Sample size (N) for ATP7A: 20, ATP7B: 13). **(E)** Colocalization quantification (Manders Colocalization Coefficient) of transfected mKO2-HA-ATP7A and eGFP-ATP7B with TGN marker p230 (Fixed cells treated with 2.5µM CuCl_2_ for 30mins) reveals their equal sensitivity to copper. Scale bar, 5µm.

To study the copper-induced trafficking routes and regulation of the two homologous copper ATPases, we expressed tagged mKO2-HA-ATP7A and eGFP-ATP7B in MDCK-II cells. Cells were allowed to completely polarize for 3-4 days on permeable filters. We found that at copper chelated and basal copper levels, both proteins localize at the TGN marked by Golgin97. Upon copper treatment (50µM CuCl_2_ for 2h) ATP7A traffics to the basolateral surface marked by ATP1A1 (Na/K-ATPase), while ATP7B localizes to the apical surface marked by gp135 (Fig. 1C). Since both proteins share high sequence similarity (∼65%) with all the conserved signature domains of copper-ATPases, we wanted to see whether they are comparably responsive to copper treatment. We analyzed their exit rate and sensitivity in terms of their trafficking from TGN. Golgi exit rate was calculated by loss of mKO2 or eGFP fluorescence from the TGN after copper treatment. We treated the cells with copper (50µM CuCl_2_) and imaged continuously at 60 sec intervals (Video1, Video2). Upon plotting the signal distribution, we found both the transporters exhibit similar Golgi-exit rate (Fig. 1D). We calculated the total fluorescence in a cell that did not change over time or on copper treatment, hence excluding the possibility of fluorescence quenching. Also, we did not observe any significant drop in fluorescence over time in untreated (control) cells expressing the fluorescent-tagged ATP7A and ATP7B, thus excluding possible quenching-related intensity drop. They showed comparable k values (i.e., ATP7A:0.03386; ATP7B:0.03225) (Fig. S1A, S1B). We also checked for the sensitivity of these proteins towards copper. We treated the cells with low copper (2.5µM CuCl_2_) to facilitate Golgi exit and measured the amount of ATP7A or ATP7B at TGN marked by p230. Calculating the Mander’s correlation coefficient we found that both Cu-exporters are stimulated to leave the TGN by identical copper concentrations (Fig. 1E, Fig. S1C, S1D). We confirmed that co-expressing ATP7A and ATP7B in polarized MDCK did not affect their respective copper-responsive localization (50µM for 2h) (Fig.S1E).

### Copper level regulates segregation of ATP7A and ATP7B into distinct TGN subdomains

ATP7A and ATP7B localize at the TGN in basal and depleted copper conditions ^57, 58^. We asked whether ATP7A and ATP7B colocalize within the TGN domains and whether copper-dependent polarized trafficking of ATP7A and ATP7B is preceded by changes in their localization within the TGN. The Copper-ATPases harbour six copper binding motifs (CXXC) on its cytosolic amino terminus and copper saturation of these sites acts as a trigger for their TGN exit ^59, 60^. We postulated that gradual saturation of the six CXXC motifs with copper might correlate with changes in the localization of ATP7A and ATP7B within the TGN, that precede their TGN exit via separate routes. To study this point, we varied the copper levels in the cell. We used *(a)* basal, *(b)* extracellular copper chelation by BCS to arrest copper uptake (25µM BCS, 4h) and *(c)* intracellular copper depletion by TTM (10µM TTM, 4h) to mimic basal copper, mild and severe copper chelation respectively in the cell.

We found that under basal copper levels, ATP7A and ATP7B localize into distinct TGN subdomains without any signal overlap. Interestingly, upon BCS treated cells we found ATP7A and ATP7B to colocalize. But surprisingly upon severe copper depletion by treatment with intracellular copper chelator, we found both the proteins at distinct non-colocalizing positions on the TGN (Fig. 2A, 2B). This indicates that the slightest change in copper concentration can dictate the intra-TGN localization of ATP7A and ATP7B. Further, our findings suggest that the two copper-ATPases colocalize at a sub-basal copper condition and are uniquely located on distinct TGN domains in condition of severe copper chelation and basal copper levels. The schematic (Fig. 2C) illustrates our findings. We confirmed the three conditions of basal copper, mild copper chelation (+BCS) and severe copper chelation (+TTM) by measuring the concentration by Inductively coupled plasma mass spectrometry (ICP-MS) (Fig. 2D).

**Figure 2.**
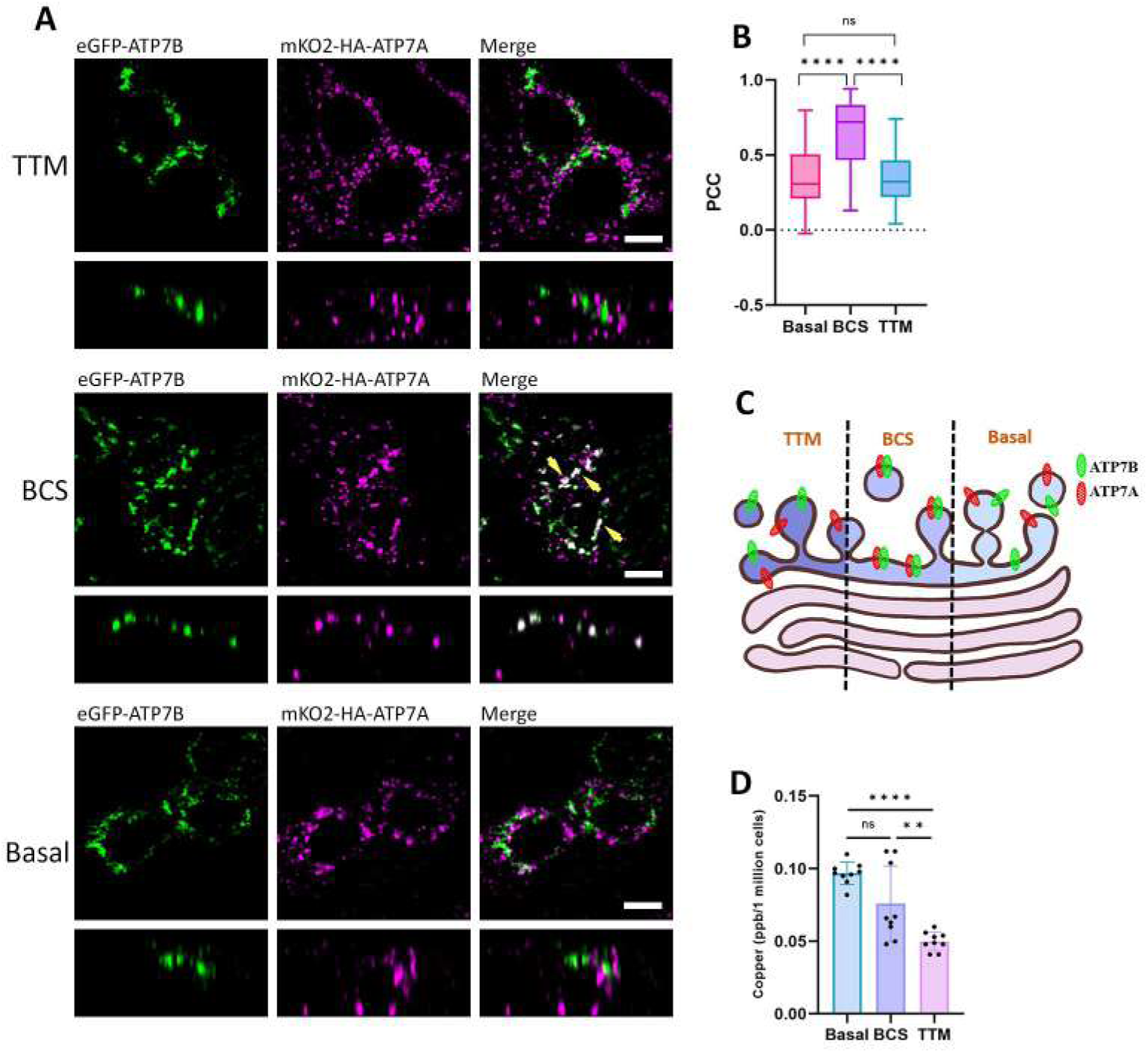
ATP7A and ATP7B occupy distinct domains at TGN under varying copper levels. **(A)** Polarized MDCK cells showing localization of eGFP-ATP7B (green) and mKO2-HA-ATP7A (magenta) under limiting copper conditions created by treatment of TTM and BCS compared to basal level. Both the Cu ATPases occupy distinct domains under TTM and basal conditions, unlike in BCS-treated cells where they colocalize, (indicated by yellow arrowheads). **(B)** Colocalization quantification (Pearson Correlation Coefficient) of transfected mKO2-HA-ATP7A with eGFP-ATP7B under basal condition, TTM and BCS treated condition (Sample size (N) for Basal: 41, BCS: 38, TTM: 20). **(C)** Schematic of our findings showing ATP7A (marked in red) and ATP7B (marked in green) occupying distinct and juxtaposed positions under the said conditions. **(D)** Measurement of intracellular copper level in basal, BCS and TTM treated conditions. This shows higher copper chelation in TTM-treated cells than in BCS and basal-treated cells. Scale bar: 5µm.

### Post-TGN trafficking of ATP7B but not ATP7A involves intermediate endosomal compartments

Extensive studies on polarized protein trafficking in MDCK cells have identified endosomal compartments that are traversed by plasma membrane targeting proteins after leaving the TGN. These compartments contribute to their apical-basolateral sorting of the trafficking proteins. In addition to their roles in endocytic recycling, Common Recycling Endosomes (CRE), Apical Recycling Endosomes (ARE), Apical and Basolateral Sorting Endosomes (ASE, BSE) have been implicated as post-TGN trafficking and sorting compartments ^28–32, 61–63^. We determined the copper-induced post-TGN trafficking itineraries of copper ATPases by studying their colocalization with markers of CRE, ARE and SE.

We treated polarized MDCK cells expressing eGFP-ATP7B or mKO2-HA-ATP7A with copper (50µM CuCl_2_) and fixed them 30 min later. We found that after TGN exit, ATP7B colocalizes with CRE, marked by transferrin (Tf) internalized for 15 mins (Fig. 3A). We also found ATP7B in Rab11 and EEA1 positive compartments, suggesting the apical route of ATP7B involves both ARE (marked by Rab11) (slow recycling) and ASE-marked by EEA1 (fast recycling) compartments respectively (Fig. 3A). In contrast with a report by Nyasae et al, we did not observe ATP7B in BSE (5 min of Tf internalization) (Fig. S2A) ^20^.

**Figure 3.**
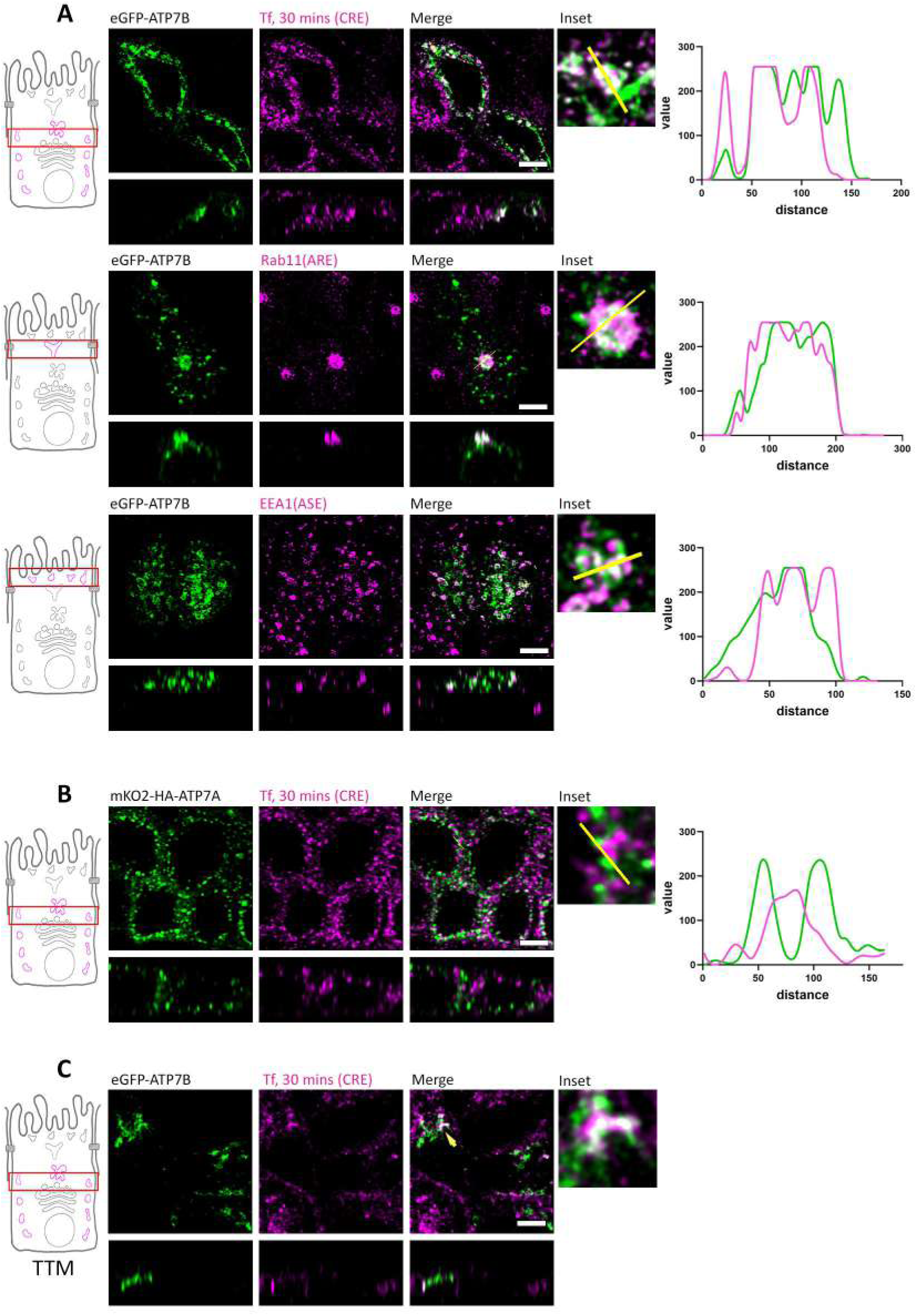
ATP7B traffic via different compartments while ATP7A does not traverse intermediate compartment. **(A)** Polarized MDCK cells showing localization of transfected eGFP-ATP7B with various intermediate endosomes as it traffics in response to copper. Insets showing zoomed images and line profiles showing pixel overlap on marked endosomes. ATP7B traffics via CRE (30mins Tf-633 internalization), ARE (Rab11 positive compartments) and ASE (EEA1 positive compartments). **(B)** Polarized MDCK cells showing localization of transfected mKO2-HA-ATP7A with CRE/BSE (30mins Tf internalization) as it traffics in response to copper. ATP7A has no overlap with CRE/BSE (30mins Tf internalization). Cells were treated with 50µM CuCl_2_ for 30 mins for (A) and (B). **(C)** Confocal image of polarized MDCK cells showing presence of eGFP-ATP7B (green) in CRE (magenta; 30 mins Tf internalization) under TTM treated condition indicating copper-independent trafficking of ATP7B (indicated by arrowhead). Scale bar: 5µm.

For ATP7A, which is targeted basolaterally, we did not observe any colocalization with CRE upon copper treatment (Fig. 3B). We also failed to observe colocalization with ASE or ARE (Rab11 and EEA1 positive compartments) (Fig. S2B, S2C). To summarize, our findings suggest that the apical route of ATP7B involves post-TGN passage through CRE, ARE and ASE whereas the basolateral route of ATP7A does not traverse any established intermediate endosomal compartments. Interestingly, we also noticed a pool of ATP7B that localizes constitutively out of TGN in basal and copper chelated condition. We determined that this ‘*constitutive out-of-TGN ATP7B*’ localized at the CRE (marked by 15 mins of TfR internalization), suggesting the presence of a copper independent recycling of ATP7B between TGN and CRE (Fig. 3C). We could not verify this phenomenon for ATP7A as we could not detect any intermediate trafficking compartment(s) for ATP7A.

### ATP7A and ATP7B exhibit both distinct and similar protein-protein interactions

Next, we sought to identify proteins that interact with ATP7B and ATP7A, thus becoming possible candidates to regulate their copper-responsive apical or basolateral trafficking. To this end, we used APEX-2-mediated proximity biotinylation ^64^; APEX-2 is an engineered peroxidase that can be rapidly induced to tag proteins quickly and efficiently with biotin phenol (BP) and H_2_O_2_ ^65^. APEX-2 tagged ATP7A and ATP7B constructs were generated (Fig. 4A) and stably expressed in MDCK cells. We confirmed that the APEX-2 constructs of ATP7A and ATP7B behave like the endogenous proteins by localizing at the TGN in basal and copper chelated conditions and trafficking to their respective plasma membrane domains upon copper treatment. Upon biotinylating the interactome of APEX-2 tagged proteins, subsequent pulldown using streptavidin beads and mass spectrometric analysis of the pull-down samples, we found a number of interacting partners for both ATP7A and ATP7B. We further manually clustered the hits based on previous reports (complete list are in Dataset S1 for ATP7A, Dataset S2 for ATP7B), to identify proteins that might be responsible for post TGN trafficking and recycling (Fig. 4B). Among a spectrum of interacting proteins, we identified various subunits of the adapter protein AP-1 complex, σ (AP1S1), β(AP1B1), µ (AP1M1, AP1M2) and Ɣ. In the present study we focused on determining the role of AP-1, as this clathrin adaptor has been previously reported to mediate apical-basolateral cargo sorting at the TGN. We confirmed our mass-spectrometric finding by co-immunoprecipitation assay where we detected AP-1 in ATP7A as well as ATP7B complexes (Fig. 4C). AP-1 has been shown to interact with ATP7A and ATP7B via the conserved C-terminal di-leucine motifs in non-epithelial models and in neurons ^50, 52^. The MEDNIK syndrome (acronym for <u>m</u>ental retardation, <u>e</u>nteropathy, <u>d</u>eafness, <u>n</u>europathy, ichthyosis, <u>k</u>eratodermia), characterized by overlapping symptoms of Wilson and Menkes disease, is caused by a mutation in the *AP1S1* gene that codes for the σ1A subunit of AP-1 ^66^. We studied the regulation of Cu-ATPases by AP-1, as epithelial cells like MDCK express 2 isoforms of AP-1 i.e., AP-1A and AP-1B ^67^. Previously, AP-1B and AP1A have been shown to play complementary roles in the sorting of basolateral cargos from TGN and endosomes ^22, 30, 68^.

**Figure 4.**
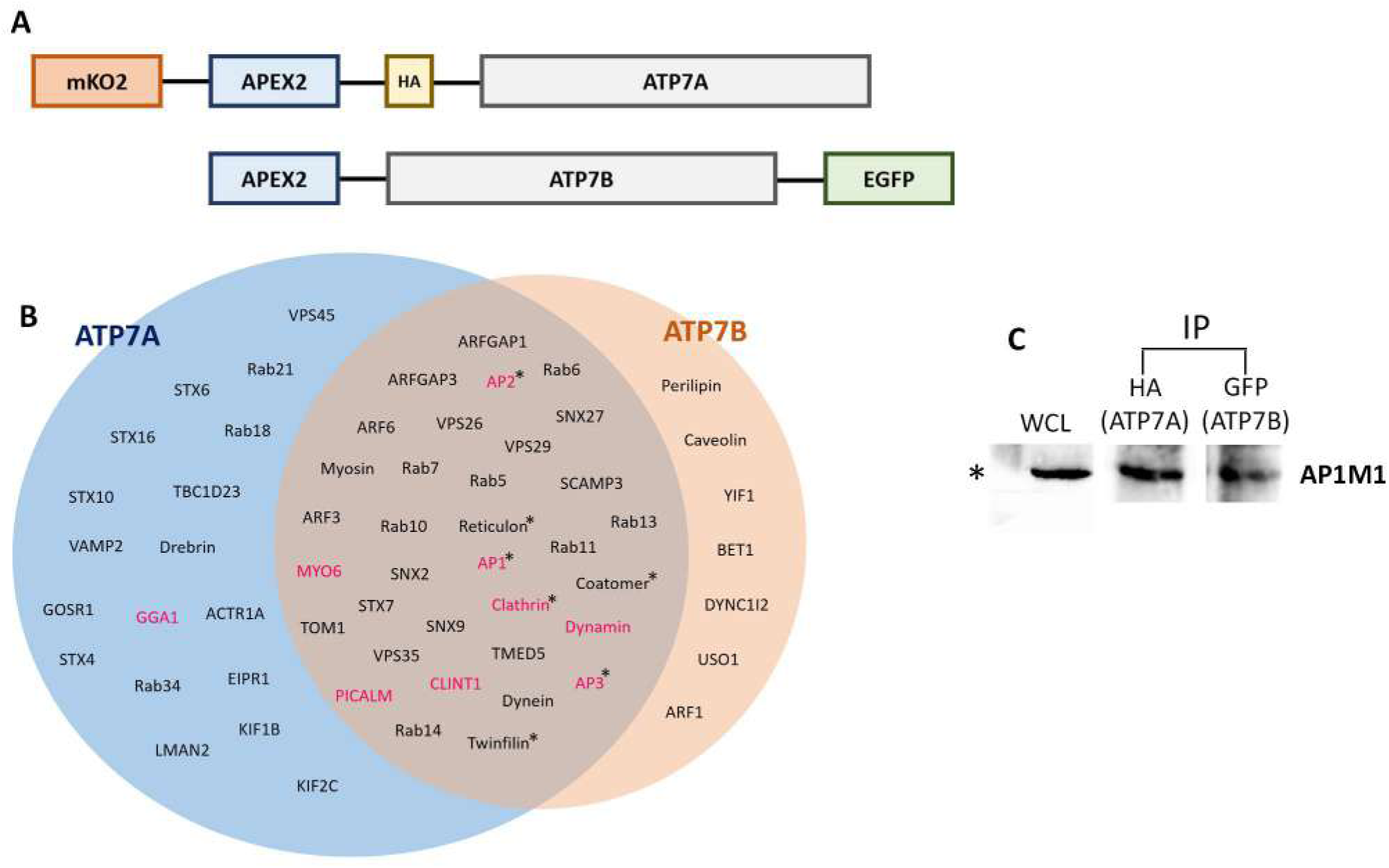
Interacting partners of ATP7A and ATP7B. **(A)** Schematic showing APEX2 tagged ATP7A and ATP7B constructs. **(B)** Venn diagram showing pulldown proteins of APEX2 tagged ATP7A and ATP7B by proximity biotinylation assay. Trafficking regulators are sorted based on previous reports. Proteins marked in magenta are putative trafficking regulators at TGN or post-TGN compartments. The ‘**\***’ mark indicates multiple subunits of those proteins are detected. **(C)** Immuno-pulldown of mKO2-HA-ATP7A and GFP-ATP7B and probed against AP-1 using rabbit anti-AP1M1 antibody, indicates interaction of AP-1 with both the copper ATPases.

### AP-1 is required for TGN retention of ATP7A and ATP7B

The clathrin adaptor AP-1 has been shown to play a crucial role in the generation and maintenance of plasma membrane protein polarity in epithelial cells. AP-1A and AP-1B isoforms differ in the possession of different μ subunits, μ1A and μ1B, respectively ^40, 67^ and in that they regulate basolateral sorting in different compartments ^68^. Albeit, it was also proposed that they have an affinity for different basolateral signals ^69^. It was initially reported that pan AP-1 knockout resulted in decrease or loss of polarization for various plasma membrane and recycling basolateral proteins such as LDLR, TfR, VSVG, ICAM1, and GPRC5A ^41, 70^, but not of apical proteins such as gp135 and non-recycling basolateral proteins such as Na/K ATPase ^42, 71^.

To investigate the roles of AP-1 in trafficking of Cu-ATPases we first generated MDCK cell lines with double knockout (KO) of AP-1A and AP-1B by knocking out the subunits AP1M1 and AP1M2. MDCK cells with pan AP-1 knockout exhibited normal growth and structural polarity comparable to wild-type cells. Notably, AP-1 KO cells exhibited higher copper accumulation in basal as well as in copper treated cells (10µM CuCl_2_; 4h), relative to wild-type cells, indicating that AP-1 plays an important role in regulating copper homeostasis in polarized epithelial cells. (Fig. 5A). This observation is in concurrence with phenotypes for MEDNIK syndrome, where mutations in AP-1A leads to imbalance in copper levels in the cell ^72^.

**Figure 5.**
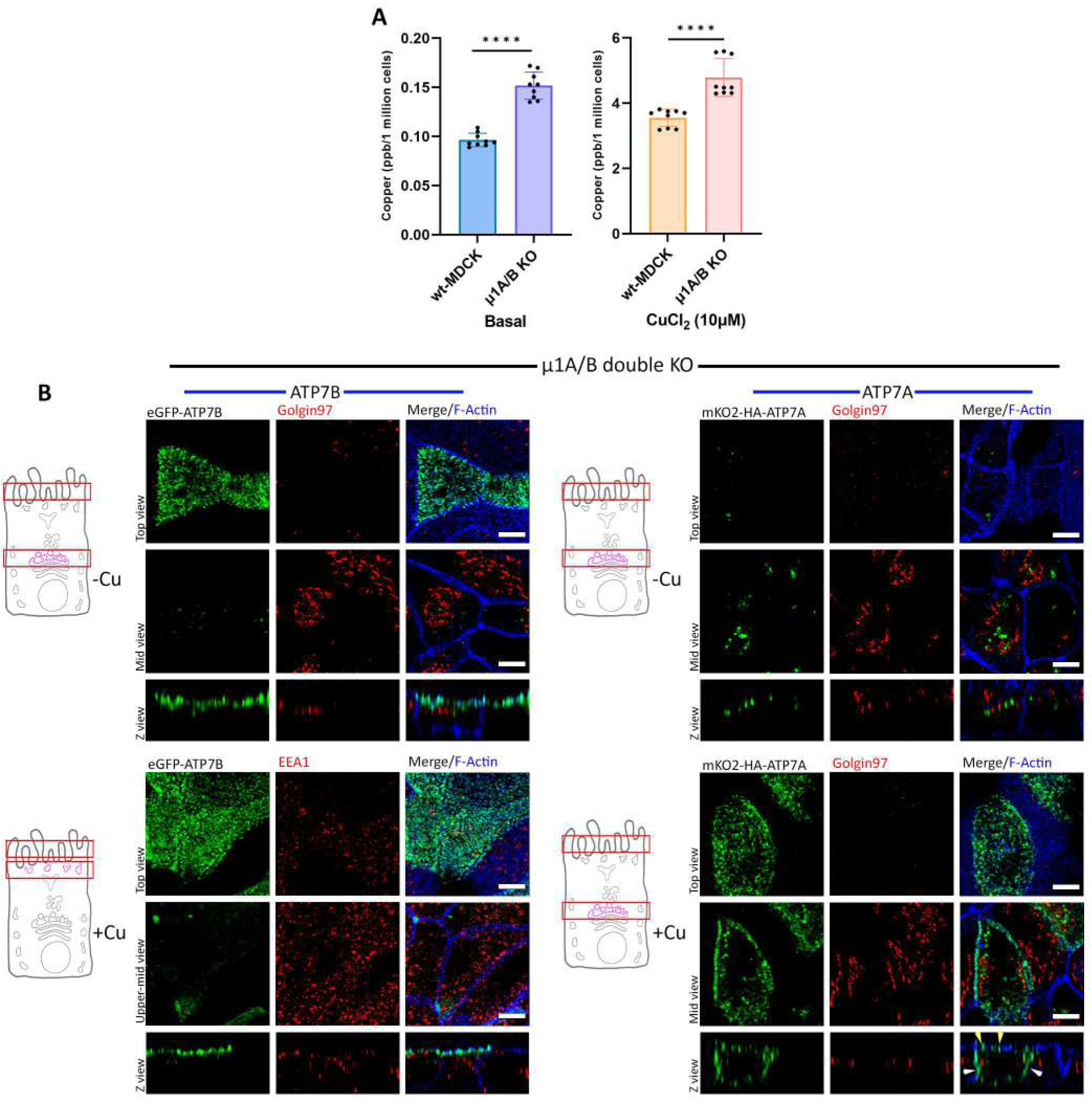
AP-1 is crucial for TGN retention of ATP7A and ATP7B. **(A)** Intracellular copper level in wild-type and AP-1 KO cells under basal as well as copper-treated conditions. AP-1 KO cells show higher copper levels in both conditions i.e., basal and CuCl_2_ treated cells (10µM CuCl_2_ for 4hrs) compared to wild-type cells. **(B)** Polarized AP-1 KO MDCK cells (µ1A and µ1B double knockout cells) showing localization of transfected mKO2-HA-ATP7A and eGFP-ATP7B. ATP7B has lost its TGN retention and constitutively localizes to the apical surface retaining its polarity in copper depleted as well as copper-treated conditions. ATP7A has also lost its TGN localization and was found to be vesicularised in copper chelated environment, but in copper-treated media it localizes to both the apical (yellow arrowhead) as well as basolateral surface (white arrowhead), losing its polarity. Scale bar: 5µm.

Pan AP-1 KO had dramatic effects on the localization of both transporters. Under copper chelated conditions both ATP7A and ATP7B lost their TGN localization, ATP7B was redistributed to the apical surface whereas ATP7A localized into a vesicular compartment. Upon copper treatment ATP7B remained on the apical surface whereas ATP7A was localized on both apical and basolateral surfaces (Fig. 5B). These experiments demonstrate that AP-1 plays a distinct role in the retention of both transporters at the TGN and in the polarized trafficking of ATP7A. Previous study on HEK293T (non-polarized cells) has shown the involvement of AP-1 in ATP7A. The protein lost its TGN localization and constitutively trafficked to plasma membrane ^50^. Interestingly our study showed that AP-1 not only regulates TGN retention but also ensures correct trafficking polarity of ATP7A, guiding it to the basolateral membrane.

### AP-1A and AP-1B cooperation regulates Copper-induced TGN exit and sorting of ATP7A and ATP7B

Next, we dissected the roles of AP-1A and AP-1B in polarized trafficking of ATP7A and ATP7B by generating MDCK cell lines deficient on either μ1A or μ1B, respectively.

In *AP-1B KO* cells, trafficking of ATP7A and ATP7B was similar to wild-type cells. At low copper levels they localized at the TGN and upon copper treatment, they trafficked to the basolateral or apical surfaces, respectively (Fig. S3). In contrast, *AP-1A KO* cells displayed a distinct phenotype. Both ATP7A and ATP7B lost their TGN localization but did not traffic to the plasma membrane even upon copper addition. Rather they localized to a vesicular compartment in both cases (Fig. 6A), unlike pan-AP-1 knockout, where they trafficked to the cell surface, albeit in a non-polarized fashion for ATP7A. Interestingly, in AP-1A KO cells, the vesicularized ATP7A and ATP7B localized at different compartments with minimal overlap. There was no change in this particular localization pattern under the effect of copper (Fig. 6B, 6C). This observation further substantiates our findings that the decision to follow two distinct itineraries by ATP7A and ATP7B happens at the TGN. Divergence towards their respective apical or basolateral pathways occurs at the TGN and not at any post-TGN compartment(s).

**Figure 6.**
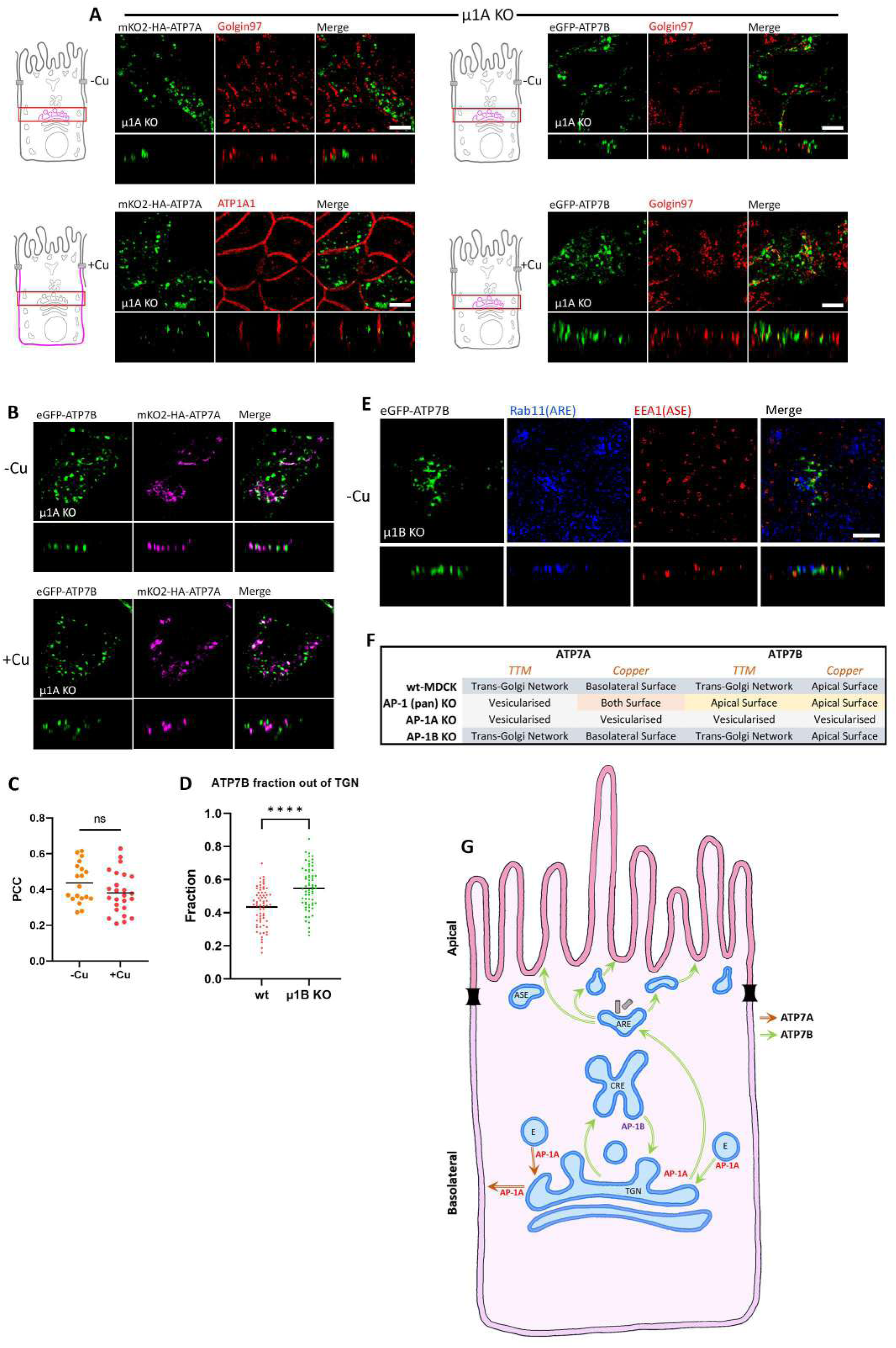
AP-1A is crucial for TGN retention as well as copper-mediated trafficking. **(A)** Images showing localization of ATP7A and ATP7B in polarized AP-1A KO MDCK cells (i.e., µ1A KO cells) in copper-deprived and elevated levels. Both the Cu ATPases lost their TGN localization and were found vesicularised in either of the copper levels and failed to reach the plasma membrane, which indicates the crucial role of AP-1A in copper-mediated trafficking. **(B)** Images showing localization of mKO2-HA-ATP7A and eGFP-ATP7B co-transfected in polarized AP-1A KO MDCK cells (i.e., µ1A KO cells) in copper-deprived and elevated levels. Both ATP7A and ATP7B localized to different endosomal compartments with minimal overlap. **(C)** Colocalization quantification (Pearson Correlation Coefficient) of transfected mKO2-HA-ATP7A with eGFP-ATP7B from (B). This shows that there is no effect of copper on the localization of ATP7A and ATP7B in AP-1A KO cells. **(D)** Fraction of transfected eGFP-ATP7B out of TGN in TTM treated cells (1 – Manders’ Colocalization Coefficient of ATP7B with Golgin97 in wild-type and AP-1B KO cells). An increased fraction of ATP7B in µ1B KO cells than wild type shows the role of AP-1B in copper-independent recycling. **(E)** Confocal images showing ATP7B in polarized AP-1B KO MDCK cells under copper chelated condition. Absence of ATP7B (green) in Rab11 (blue) and EEA1 (red) positive compartments suggests lack of anterograde trafficking or spillover from CRE due to regulation by AP-1B in basolateral trafficking at CRE. **(F)** Tabulated summary of trafficking phenotypes of ATP7A and ATP7B in AP-1A KO, AP-1B KO and AP-1 pan KO cells. **(G)** Proposed model for copper-independent and copper-dependent trafficking of both the Copper ATPases, i.e., ATP7A and ATP7B, in polarized MDCK cells. Common endosomal stations are marked like TGN (Trans-Golgi Network), CRE (Common Recycling Endosome), ARE (Apical Recycling Endosome), ASE (Apical Sorting Endosome) and E (Endosome). Red arrow marks the trafficking itinerary of ATP7A and the Green arrow represents the trafficking path of ATP7B. Scale bar: 5µm.

Our results show that AP-1A functions at TGN providing trafficking directionality and TGN retention for both ATP7A and ATP7B. On the other hand, AP-1B does not play a detectable role in the presence of AP-1A, but in the absence of AP-1A, AP-1B provides a back-up mechanism to facilitate transport of both transporters to the cell surface, though in a non-polarized fashion for ATP7A. Interestingly, ATP7B, but not ATP7A, also localizes at the CRE (marked by 15 mins pulse-chase internalization of TfR) in copper-chelated or basal conditions (Fig. 3C), suggesting Cu-independent trafficking of ATP7B from TGN and CRE and, possibly, AP-1B-dependent return from CRE to TGN. Consistently, AP-1B KO resulted in a significantly higher pool of ATP7B dispersed out of TGN in copper-chelated condition (Fig. 6D). AP-1B KO does not facilitate ATP7B’s trafficking from CRE to ARE and/or ASE under copper-chelated condition (Fig. 6E). In contrast, Perez-Bay and co-workers showed that in AP-1B knocked-down MDCK cells, TfR further mislocalizes to ARE deviating from its original PM-BSE-CRE basolateral itinerary ^45^.

These observations reveal an additional role for AP-1B in ATP7B’s (copper-independent) anterograde route towards the apical plasma membrane. In contrast, copper-dependent trafficking of ATP7B does not involve its CRE localization or regulation by AP-1B. The phenotypes that we observed for ATP7A and ATP7B in the knockout cells in different copper conditions have been summarized in (Fig. 6F). Based on our observations in the three KO cell lines, we propose a model that illustrates the roles of AP-1 in sorting and trafficking of ATP7A and ATP7B (Fig. 6G).

### ATP7B mutations suggest additional mechanisms for its TGN retention and subsequent polarized sorting at the TGN

Despite sharing high sequence homology, ATP7B differs completely from ATP7A in its proximal amino-terminus domain (amino acids 1-63) (Guo et al., 2005), which is critical for TGN exit and proper apical sorting in hepatocytes and WIF-B cells (Hasan et al., 2012). A stretch of 9 amino acids (F^37^AFDNVGYE^45^) forms the core minimum motif, sufficient for ATP7B apical sorting even in copper limiting conditions. It contains a ^41^NVGY^44^ putative Clathrin-binding motif and a N41S hot-spot mutation responsible for Wilson Disease ^47^.

We observed that deletion of either 9 residues (ΔF^37^-E^45^) or the core ^41^NVGY^44^ motif resulted in ATP7B basolateral sorting under copper stimulation (Fig.7A) and faster Golgi exit rate than the wt-ATP7B (k value 0.2248 vs 0.03225, Fig.7B, S4B and Video3), which is a reminiscent, albeit milder, phenotype of ATP7B losing its TGN retention even in copper limiting condition in AP-1A KO cells (Fig 6A, top image, right panel). Similar results were observed in the ^41^NVGY^44^ deletion and N41S mutation (Fig. S4A).

**Figure 7.**
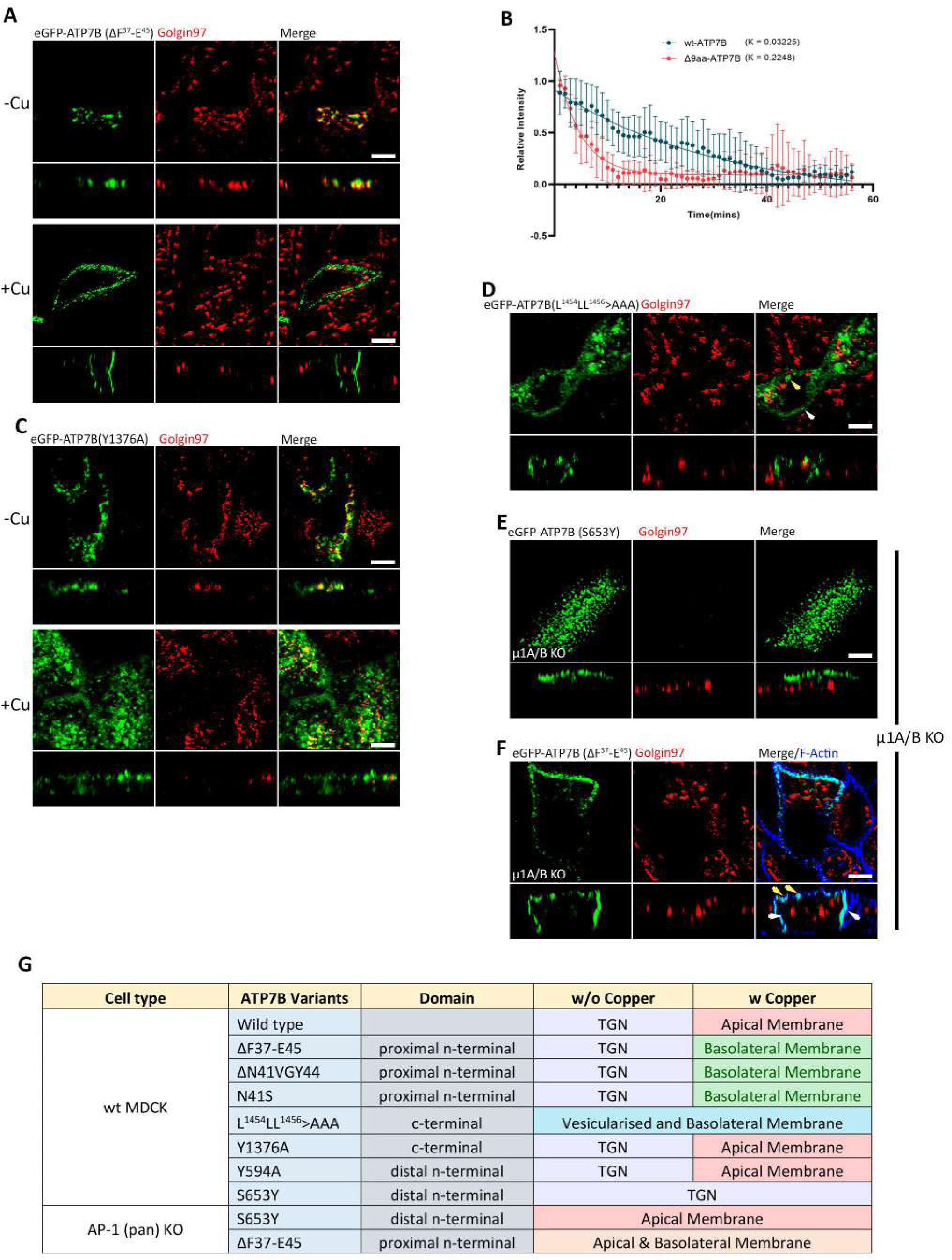
ATP7B mutants reveal additional regulation for Apical polarity. **(A**) Images showing 9 aa deletion mutant (ΔF^37^-E^45^-ATP7B) trafficking to basolateral surface in copper treated condition. **(B)** Fitted curve of decreased pixel count of eGFP-ATP7B and Δ9aa mutant in copper treated condition showing dispersion of Cu ATPases and their exit rate. Higher K value of Δ9aa mutant shows a faster Golgi exit rate than the wild type ATP7B (Sample size (N) for ATP7B: 13, Δ9aa mutant: 6). **(C)** Confocal image of Y1376A mutant of ATP7B showing wild-type-like phenotype. **(D)** Image showing tri-leucine mutant of ATP7B (i.e., L^1454^LL^1456^>AAA) has lost its TGN localization and localized to the basolateral surface (white arrowhead) as well as in endosomes (yellow arrowhead). **(E)** Image of S653Y (a non-Golgi exiting mutant of ATP7B) in AP1 KO cells (i.e., µ1A/B double knockout cells) informs its constitutive localization on the apical surface of the plasma membrane. **(F)** Confocal image of ΔF^37^-E^45^ (a basolateral targeting mutant) in AP1 double KO cells show loss of TGN retention as well as in its polarity because of constitutive localization to both the surfaces of the plasma membrane (apical surface marked by yellow arrowhead and basolateral surface marked by white arrowhead). Scale bar, 5µm.

Since previous studies on dileucine in the carboxy-terminal of ATP7B have shown its AP-1 interaction and its involvement in somatodendritic polarity ^52^, we screened the cytosolic domains of ATP7B for the presence of additional Clathrin-interacting motives. Mutations on two YxxФ motives on the C-term (Y^1376^>A. Fig. 7C) or the N-Term (Y^594^A, Fig. S4C) showed wild-type trafficking phenotype. Mutating the tri-leucine motif (L^1454^ –L^1456^) essential for retrograde trafficking ^73^ (L^1454^LL^1456^>AAA) resulted in loss of TGN localization, even in TTM-treated cells (Fig. 7D), similar to the phenotype observed in AP-1A KO cells.

A Copper-insensitive, Wilson Disease-causing mutant ATP7B-S653Y ^62^ did not exit TGN in WT MDCK (Fig. S4D), but trafficked to the apical plasma membrane in a Cu-independent manner in pan-AP-1 KO cells (Fig. 7E). On the other hand, ATP7B-ΔF^37^-E^45^ mutant in pan-AP-1 KO cells also lost its TGN retention in copper limiting condition, that was originally preserved in wild-type cells.

Interestingly it loses its polarity to both the membrane surfaces contrary to the wild type where it was localized only to the basolateral surface. (Fig. 7F). These findings substantiate the regulatory role of AP-1 in TGN retention of the Copper-ATPases. The phenotypes that we observed for ATP7B mutants in the wild-type and knockout cells in different copper conditions have been summarized in (Fig. 7G).

To summarize, our studies on ATP7B mutants expressed in wild-type and AP-1 KO cells suggest a strong role of AP-1 in TGN retention, but the directionality of ATP7B depends on AP-1 in synergy with other regulatory signals or regulators. Mutating putative Clathrin binding motifs in ATP7B failed to replicate the exact itinerary observed in AP-1 KO cells. This indicates the presence of a multitier regulatory mechanism of copper-dependent trafficking of ATP7B. Further studies on this regulation can be fascinating as polarized epithelial cells can reveal new insights into stepwise control of polarized trafficking.

Our study sheds light on the molecular mechanism of overlap of phenotypes of copper metabolism diseases, Wilson and Menkes disease with MEDNIK syndrome. Our study also unravels the presence of a default trafficking pathway that directs the protein to the basolateral membrane.

## Discussion

Copper levels in a mammalian cell is primarily regulated by the homologous P-type copper-ATPases, ATP7A and ATP7B. We determined the underlying mechanism of basolateral vs apical trafficking of ATP7A and ATP7B respectively in response to elevated copper in a model of polarized epithelia. ATP7A and ATP7B were first determined as two distinct copper-transporting ATPases in fish. In invertebrates, possibly a single copper-ATPase was responsible for copper transport in the biosynthetic pathway as well as for exporting excess copper. With the evolution of two copper ATPases, the functional responsibility was divided with respect to (a) copper uptake from the alimentary canal to the bloodstream for systemic copper distribution and (b) biliary copper export for detoxification. In parallel to this functional specialization, the trafficking polarity of the two copper ATPases came into existence, with copper-distributing ATPase, ATP7A, which transports copper into the bloodstream attaining basolateral trafficking polarity and the copper-detoxifying ATPase, ATP7B, gaining the apical or luminal trafficking polarity. We asked what could be the possible mechanism that regulates the opposite trafficking polarity of such evolutionary and functionally similar membrane transporters.

Epithelial cells contain two isoforms of AP-1 clathrin adaptor complexes. AP-1A is ubiquitously expressed and regulates transport between the TGN and endosomes. AP-1B is expressed only in epithelia and mediates the polarized targeting of membrane proteins to the basolateral surface ^74^. A study from Bonifacino’s group was the first to identify a novel member of the adaptor medium chain family, μ1B, which is closely related to μ1A and shares a 79% amino acid sequence identity ^40, 67^. In a review that summarized the developments in the field of Clathrin over the last 40 years, Robinson mentioned that Vertebrates were the first organisms where epithelial cell-specific isoforms of the AP-1µ subunit emerged, which contributes to basolateral sorting ^75^. We hypothesize that the divergence of homologous membrane proteins towards the apical and basolateral membrane of polarized epithelia coincided with appearing of the two isoforms of AP1, i.e., AP-1A and AP-1B.

Using a mass-spectrometric approach we identified multiple partners that could regulate the apico-basolateral trafficking polarity of ATP7A and ATP7B. A pool of hits emerged that constituted subunits of Adaptor proteins (APs), specifically, AP1. The AP complexes are heterotetrameric protein complexes that mediate intracellular membrane trafficking at endocytic and secretory pathways. There are five different AP complexes: AP-1, AP-2 and AP-3 are Clathrin-associated complexes; whereas AP-4 and AP-5 are not ^76^. Mutations on AP subunits have been implicated in a variety of inherited disorders, that includes: MEDNIK (mental retardation, enteropathy, deafness, peripheral neuropathy, ichthyosis and keratodermia) syndrome, Hermansky–Pudlak syndrome, Fried syndrome, and HSP (hereditary spastic paraplegia) ^77–80^. AP-1 is a highly conserved Clathrin adaptor that functions primarily at the trans-Golgi network (TGN) and also at endosomes and lysosomes ^81^. Growing evidences have implicated the involvement of AP-1 in disorders of copper metabolism. MEDNIK symptoms caused due to mutation in AP1-σ shares multiple symptoms and cellular phenotypes as of Wilson and Menkes disease, hence indicating the possible involvement of AP-1 in regulation of ATP7A and ATP7B ^66^. Recently, Alsaif et al identified mutations in AP1B1 that encode the large β subunit of the AP-1 complex causing MEDNIK-like syndrome ^78^. The affected individuals manifested abnormal copper metabolism, evidenced by low plasma copper and ceruloplasmin, but normal levels of hepatic copper. We also detected higher copper in AP-1 knockout cells. Our mass-spectrometric data and these previous genetic and clinical reports prompted us to conduct an indepth study of the regulation of the Wilson and Menkes disease proteins by the Adaptor Protein, AP-1. Though it has been well established that in polarized epithelial cells, ATP7A traffics to the basolateral end and ATP7B towards the apical upon copper treatment, the trafficking itinerary in these cells has not been determined. We found that ATP7A follows a direct route from the TGN that bypasses any known sorting compartments that have been implicated in basolateral trafficking, e.g., CRE, basolateral sorting or the basolateral endosomes. On the other hand, ATP7B traverses through a more elaborate itinerary that involves CRE, ASE and ARE. Interestingly, our finding shows contradiction to Nyasae et al.’s findings in WIF-B cells, where ATP7B localizes at the BSE before being transported to the apical membrane. Differences in cell type and lineage may be responsible for this contrasting finding.

We discovered a pool of ATP7B (and not ATP7A) that exhibits copper-independent post-TGN and CRE localization. Interestingly, this pool of copper-independent ATP7B is distinct from the more abundant copper-exporting ATP7B pool that traverses through the ARE and ASE. We hypothesize that these compartments may serve as a storage site for copper that might be eventually utilized in situations of copper deficiency. However further investigation is warranted to establish this model. We found that AP1B regulates this distribution of ATP7B between TGN and the CRE.

To summarize, we propose a model that illustrates the roles of AP-1 in sorting and trafficking of the homologous copper transporting ATPases, ATP7A and ATP7B. AP-1A functions at TGN providing directionality and TGN retention for both ATP7A and ATP7B. In absence of AP1A, AP1B works at the TGN-retention checkpoint for both the Cu-ATPases. This finding is in agreement with Gravotta et al ^42^ where they demonstrate a critical role of the ubiquitous AP-1A in basolateral sorting. Knockdown of AP-1A causes missorting of basolateral proteins in MDCK cells, but only after knockdown of AP-1B, suggesting that AP-1B can compensate for lack of AP-1A.

In concurrence with our findings, they also suggest that knockdown of AP-1B promotes ‘spillover’ of basolaterally trafficking proteins into CRE, the site of function of AP-1B, suggesting complementary roles of both adaptors in basolateral sorting. We found that in absence of both AP-1A and AP-1B, ATP7A loses its trafficking polarity and constitutively localizes at both membranes. On the other hand, ATP7B loses its TGN retention capacity but retains trafficking polarity that now becomes independent of intracellular copper concentration.

The importance of our study lies in establishing the roles and understanding the mechanism of the Clathrin adaptor protein AP-1 in copper metabolism by regulating the trafficking dynamics of the Wilson and Menkes disease proteins. Many patients exhibit a spectrum of Wilson disease-like clinical symptoms, but upon gene sequencing, *ATP7B* or copper-metabolism pathways genes reveal no mutations. It will be important to screen AP-1A and AP-1B in those patients that might reveal novel mutations thus enabling better disease diagnosis.

## Materials and Methods

### Plasmids and Antibodies

eGFP-ATP7B construct was available in the lab previously^82^. HA-ATP7A was a kind gift from from Michael Petris (University of Missouri) and Svetlana Lutsenko (Johns Hopkins University). mKO2 fluorescent tag was added to the N-terminal of ATP7A with the final construct as mKO2-HA-ATP7A. APEX2-ATP7B-EGFP and APEX2-mKO2-HA-ATP7A constructs were made on the existing plasmid using NEBuilder HiFi DNA Assembly (NEB #E2621). Mutations on eGFP-ATP7B were prepared following Q5 Site-Directed Mutagenesis Kit (NEB #E0554) protocol.

Primers used for ATP7B mutants:

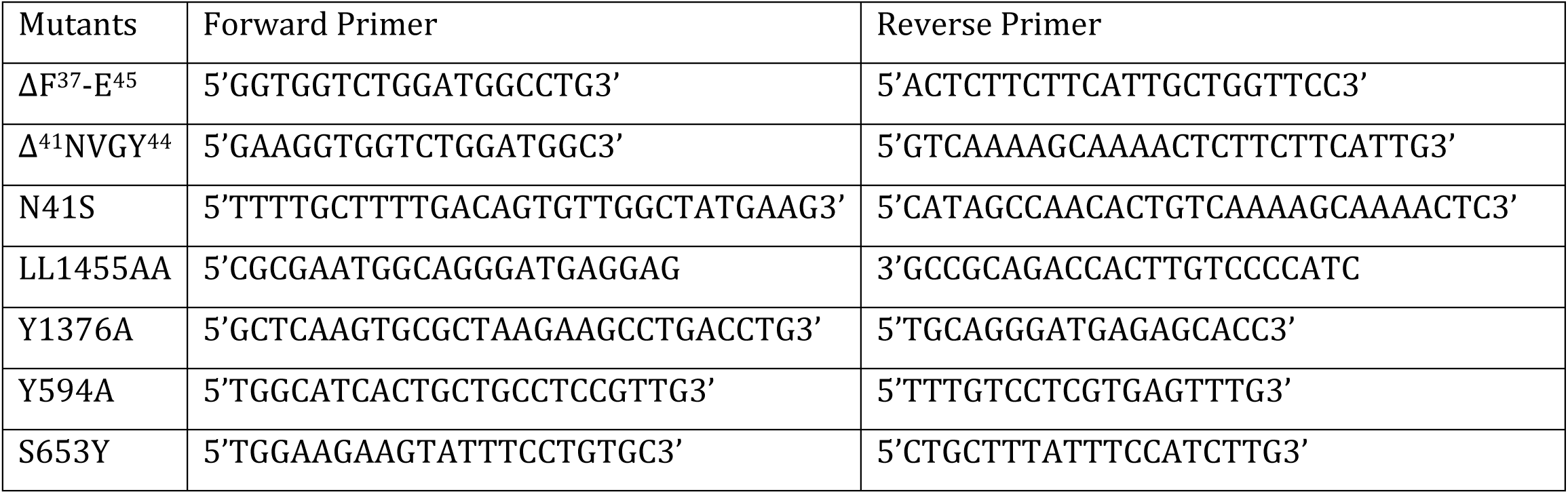

Plasmid isolation was done using Macherey Nagel plasmid isolation kit (#MN740490).

Following antibodies are used: mouse anti-Golgin97 (Thermo #A21270), mouse anti-p230 (BDBiosciences #611280), mouse anti-ATP1A1 (Abcam #ab7671), mouse anti-gp135 (DSHB #3F2/D8), mouse anti-EEA1 (BD Transduction #610456), rabbit anti-Rab11 (Thermo #715300), rabbit anti-AP1M1 (Sigma #SAB1301057), mouse anti-HA (Biolegend #901501), mouse anti-GFP (DSHB #DSHB-GFP-12A6), anti-mouse Alexa 488 (Invitrogen #A11029), anti-mouse Alexa 555 (Invitrogen #A32773), anti-rabbit Alexa 568 (Invitrogen #A11011), anti-mouse Alexa 647 (Invitrogen #32787), anti-rabbit HRP conjugated (CST #7074S).

### Cell lines and Cell culture

MDCK cells were grown and maintained in media consisting of DMEM (Gibco #11995056), supplemented with 10% FBS (Gibco #26140079) and 1X PenStrep (Gibco #15140122) under 5% CO_2_. For monolayer polarization 3 x 10^5^ cells were plated in 0.4µm, 12mm inserts (Corning #3401) and grown for 3-4 days. For transfection, electroporation was performed using Nucleofector 2b device (Lonza #AAB-1001) and Amaxa Kit V (Lonza #VCA-1003) Program T-023.

For copper treatment, 50uM of CuCl_2_ was used for 2 hrs, else otherwise mentioned. For mimicking copper-deprived conditions, cells were treated with 50uM BCS or 25uM TTM for 2hrs.

For the establishment of stable cell lines, respective plasmids are transfected and cells are maintained in selection media i.e., 1ug/mL puromycin (Gibco #A1113803) for 48 hrs. After that cells were grown in fresh media and visible colonies are isolated using cloning cylinders and subjected to Fluorescence-activated cell sorting (FACS), only selecting low to medium-expression cells.

### Immunofluorescence and microscopy

After desired treatments, cells were washed twice with ice-cold PBS, then fixed with 2% PFA for 20 mins. Cells are washed with PBS followed by 20 mins incubation in 50mM ammonium chloride. Next, the cells were washed with PBS, and blocking and permeabilisation were performed in 1% Bovine Serum Albumin (BSA, SRL #85171) in PBSS (0.075% saponin in PBS) for 20 mins. Primary antibody incubation was performed for 2 h at RT followed by three PBSS washes. After that, incubation with the respective secondary antibodies was done for 1 h followed by three PBSS washes. The membrane was mounted with the ProLong Gold Antifade Reagent (CST #9071). All images were acquired with Leica SP8 confocal platform using oil immersion 63x objective (NA 1.4) and deconvoluted using Leica Lightning software.

For the Golgi-exit rate, cells are transfected with eGFP-ATP7B or mKO2-HA-ATP7A and plated in glass bottom dishes or slides. Before imaging, phenol-containing media was replaced with imaging media (DMEM w/o phenol red (Gibco #31053028), 20mM HEPES (Gibco #15630106), 1mM Trolox(Sigma #238813), 1% FBS). 50µM CuCl_2_ was treated, and cells were imaged at an interval of 1 min.

### Image Analysis

For calculating the Golgi exit kinetics, a uniform mean-intensity range was used in all the slices by resetting the min and max intensity. From the histogram, pixels having more than 95% of the maximum intensity value (in this case 65653 as 16-bit image is used) were considered and counted as Golgi in basal condition. Upon copper treatment, the disappearance of those pixels was considered as Golgi exit. Decrease in pixel count was plotted against time. All images were processed and analyzed using ImageJ software. The Macro code for the analysis is available at https://github.com/saps018/7A-7B-.

For colocalization calculation, the JaCoP plugin was used. Mander’s or Pearson’s colocalization coefficient was calculated from manually drawn ROIs. Graphs were plotted using Graphpad Prism (version 9.4).

### Tf internalization assay

For 633-Tf (Invitrogen #T23362) uptake assays, cells were starved for 60 min at 37°C in HBSS containing 20 mM HEPES (standard buffer) and incubated for 60 min at 4°C with 15 μg/ml 633-Tf in 1% BSA standard buffer. After that, the temperature was shifted to 37°C and after the specified time, cells were immediately rinsed (with ice-cold HBSS) and fixed (ice-cold 2% PFA in PBS). After quenching PFA (with 50 mM NH_4_Cl in PBS) for 15 min, cells were permeabilized with 0.01% Triton X-100 in PBS for 7 min. Following this, the cells are incubated with the corresponding primary antibodies for 2 h and then secondary antibodies again for 2 h in 1% BSA in PBS and washed three times after each incubation.

To label BSE (basal sorting endosome) and CRE (common recycling endosome), Tf-633 was added to the basolateral side of the cells, growing on the membrane, for 5 min and 30 min, respectively, at 37°C (15 μg/ml in 1% BSA standard buffer).

### Reverse transcription and PCR

MDCK cells were seeded in 60mm culture dishes. After confluency, media was discarded, and cells were harvested using 1mL TRIzol reagent (Thermo #15596018). All the downstream processes were performed according to the manufacturer’s protocol. Briefly 0.2mL chloroform per 1mL of TRIzol was added to the harvested cells for lysis. Centrifuged for 15mins at 12000x g at 4°C. The aqueous phase containing RNA was carefully isolated and 0.5mL of isopropanol was added. Centrifuged for 10 mins at 12000x g at 4°C for precipitation of RNA. After two ethanol washes, RNA was air dried and reconstituted in Nuclease-free water and stored at -80°C. cDNA was prepared from 1µg of previously prepared RNA using Verso cDNA Synthesis Kit (Thermo #AB1453A).

Polymerase chain reaction (PCR) was performed using 1µL of cDNA and below said primers. Q5 High-Fidelity 2X Master Mix (NEB #M0492S) was used for the PCR reaction according to the manufacturer’s protocol.

Primers used for reverse transcription PCR:

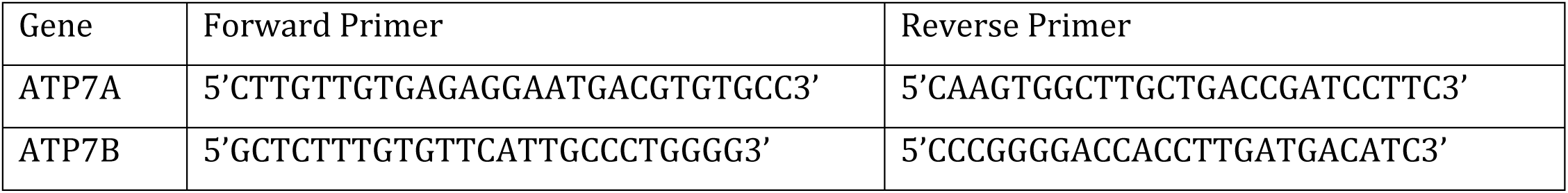

### Intracellular copper measurement

An equal number of cells (wild-type MDCK or AP-1 KO cells) were seeded in 60mm culture dishes. After they reached confluency, cells were treated with either TTM, BCS, CuCl_2_ or basal copper conditions. Cells were then washed with ice-cold PBS several times (5-6 times) and were harvested in centrifuge tubes. Next cells were counted, and an equal number of cells were digested in 100uL of Nitric acid (Merck #1.00441.1000) overnight at 95°C. Copper standards were prepared from 23 Element standard (Reagecon #ICP23A20). To bring the final concentration of nitric acid ≤2%, digested samples were diluted in 5mL Mili Q water (Millipore) and analyzed using Xseries 2 ICP-MS (Thermo Scientific). Values were plotted using GraphPad Prism.

### Proximity biotinylation

Stable cells expressing either APEX2-mKO2-HA-ATP7A or APEX2-ATP7B-GFP are plated in 0.4uM 24mm inserts and cultured till confluency. 2mM biotinyl tyramide (Sigma # SML2135) containing media was added 30 mins prior to Copper treatment. Cells were washed in PBS^++^ (with Ca^2+^/Mg^2+^) three times. The peroxidase reaction was performed for 1 min using 0.5mM H_2_O_2_ followed by quenching in quencher (PBS++ with 10mM Sodium ascorbate, 5mM Trolox and 10mM Sodium azide). Subsequently, cells are washed with quencher twice. Next cells are scrapped in RIPA lysis () and flash frozen. Samples are thawed and sonicated in medium amplitude for 30 secs. Lysates were clarified by centrifugation at 13000 rpm for 10 minutes at 4°C.

For the isolation of biotinylated proteins, Streptavidin-coated magnetic beads (Genscript #L00936) were prepared by washing twice with RIPA lysis. Clarified lysates from the above step were added to the beads and incubated at room temperature for 2 hours with gentle mixing. For each sample 100uL of bead slurry was used. Streptavidin beads were then washed three times with RIPA lysis. Biotinylated proteins were eluted in 1.5x NuPAGE LDS Sample buffer (Invitrogen #NP0007) containing 20mM DDT (SRL #17315) and 2mM biotin (SRL #18888) by heating at 95°C for 10 minutes.

### In-gel digestion and extraction

Biotinylated proteins eluted from magnetic streptavidin beads were separated on a NuPAGE Novex Bis-Tris 4-12% gel run for 10 minutes hour at 120V. Lanes for each sample were manually cut into 1x1 mm cubes. Gel bands were destained with 300-800 µl of 70% 50mM ammonium bicarbonate/30 % Acetonitrile for 30-60 minutes followed by dehydration with acetonitrile. Dehydrated gel bands were swelled with 150 µL of 10 mM DTT in 50 mM ammonium bicarbonate for 45 minutes at 56°C. The DTT solution was subsequently removed and replaced with 150 µL of 55 mM iodoacetamide in 50 mM ammonium bicarbonate and incubated in dark for 30 minutes. The iodoacetamide solution was removed and bands were dehydrated with acetonitrile. Approximately 30-50 µL of 15 ng/µL Sequencing Grade Trypsin (Promega) was added to each sample and digestion was completed overnight at 37°C. After the overnight digestion, excess trypsin solution was removed from each sample and bands were incubated with 150 µL of extraction solution (5% formic acid, 30 % acetonitrile). The extraction solution was collected after 10 minutes, and this process was repeated once. The final peptide extraction was completed by incubating gel bands with 100% acetonitrile for 5 minutes. Extracted peptides were dried to completeness in a vacuum concentrator. For desalting, samples were reconstituted in 20 µL of 0.1% formic acid and loaded onto C18 StageTips. The tips were washed twice with 50 µL of 0.1% formic acid, and peptides were eluted with 50% acetonitrile/0.1% formic acid and then dried in a vacuum concentrator.

### Liquid chromatography and mass-spectrometry

Samples were analysed on a Q-Exactive HF mass spectrometer coupled with nanoflow HPLC System (Easy-nLC 1200 Thermo Scientific). The sample was loaded on to Easy Spray Column PepMapTM RSLC C18 (3 μm, 100 A^0^ 75 μm x 15 cm). Samples were eluted with a 60 minutes gradient between solution A (5 % acetonitrile-water containing 0.1 % formic acid) and solution B (95 % acetonitrile in water containing 0.1% formic acid) at the flow rate of 300 nl/min as detailed below.

| Time (min) | % sol. B |
| --- | --- |
| 0 | 3 |
| 41 | 25 |
| 48 | 40 |
| 54 | 95 |
| 60 | 95 |

MS was acquired with Orbitrap analyzer at the resolving power of 60,000 at m/z 200. Scan range selected was 400-1650 m/z. MS/MS was carried out in HCD (normalized collision energy - 28%) mode with resolving power of 15,000 at m/z 200. Top 10 ions were taken for MS/MS. The lock mass option was enabled for accurate mass measurements.

Proteins and peptides were identified and quantified using the MaxQuant software package (version 2.1.0.0)^83, 84^. Data were searched against the UniProt canine database which contains 59,102 entries. A fixed modification of carbamidomethylation of cysteine and variable modifications of N-terminal protein acetylation, oxidation of methionine, and biotin-phenol on tyrosine were searched. The enzyme specificity was set to trypsin allowing cleavages N-terminal to proline and a maximum of two missed cleavages was utilized for searching. The maximum precursor ion charge state was set to 6. The precursor mass tolerance was set to 20 ppm for the first search where initial mass recalibration is completed and 4.5 ppm for the main search. The MS/MS tolerance was set to 20 ppm and the top MS/MS peaks per 100 Da was set to 12. The peptide and protein false discovery rates were set to 0.01 and the minimum peptide length was set to 6.

### Co-immunoprecipitation and immunoblot analysis

MDCK cells expressing either mKO2-HA-ATP7A or eGFP-ATP7B were seeded in equivalent numbers and grown to 60-70% confluence in 60mm culture dishes for 72 hours. On the day of the experiment, cells were treated with 50uM Copper for 15 mins subsequently which they were lysed using syringe in non-degrading lysis buffer (50mM Tris-Cl pH 7.5, 150 mM NaCl, 0.1% CHAPs, 1mM EDTA supplemented with PIC, PhosSTOP and Na_3_VO_4_) for up to 1hr. Lysates were centrifuged at 10,000 rpm at 4°C for 5 mins. The supernatant, i.e., the crude cell lysate was then subjected to Bradford Assay to measure the total protein concentration. Simultaneously, Protein G coated MagBeads (Genscript #L00274) were equilibrated with Binding/Wash Buffer (20mM Na_2_HPO_4_, 150mM NaCl, pH 7.0) and incubated with the target IgG at room temperature while mixing on a rotator for 60 minutes. Cross-linking of mouse anti-HA or mouse anti-GFP to the MagBeads was done using freshly prepared cross-linking solution (20 mM dimethyl pimelimidate dihydrochloride in 0.2 M triethanolamine, pH 8.2) for 30 minutes at room temperature using rotational mixing. The cross-linking reaction was stopped using 50 mM Tris, pH 7.5 for 15 minutes at room temperature using rotational mixing post which the cross-linked beads were washed three times with PBS, pH 7.4. An equal amount of protein extracts (100ug) were added to the cross-linked MagBeads for overnight binding in a rotator at 4°C. After 3 washes of the protein-captured beads with PBS, 1X LDS Sample Buffer was added and boiled at 95°C for 10 min to denature the proteins and separate them from the beads. The proteins were separated in SDS-PAGE and transferred onto PVDF membranes. After blocking with 3% BSA in TBST for 2hrs, the membranes were probed with primary antibodies against AP1M1 at 4°C overnight, followed by secondary antibody conjugated to HRP. Then, the membranes were developed (Bio-Rad #1705062) and visualized using ChemiDoc (Bio-Rad).

### Generation AP1 knockout MDCK-II cell lines

For AP1 subunit genetic knockout, the CRISPR Cas9 mediated non-homologous end joining (NHEJ) strategy was implemented. Briefly, guideRNA selection was determined by picking the first common exon based on the NCBI Gene curated transcript database for each gene. The first shared exon of all known transcripts was then used as the “target” for guide selection in the CHOPCHOP guide selection tool (http://chopchop.cbu.uib.no). The highest scoring 19bp guide was then synthesized with an additional guanidine nucleotide at the 5’ end of the guide sequence for efficient U6 promoter transcription, and with BbsI sticky-ends for cloning into the BbsI linearized pX459v2 ^85^. The guide sequences used were: AP1M1 - TCATCTGCCGGAATTACCG; AP1M2 - CATGCCTCTGCTCATGCAG.

4 million MDCK cells were transfected with 5 micrograms of the pX459v2-AP1M1 or pX459v2-AP1M2 or both the plasmids with the Lonza AMAXA 2B nucleofection system. 24 hours after plating the freshly transfected cells to 150mm dishes (30,000 and 1,000 cells per dish) 1µg/mL puromycin was added and after 2 days fresh media is added. After an additional 7 to 10 days to expand the single clones, glass cloning cylinders were used to trypsinize and plate the clones to a 96 well plate. These clones were sequence verified to select knockout of the desired gene.

## Acknowledgements

This work is supported by DBT-Wellcome Trust India Alliance Fellowship (IA/I/16/1/502369), and Core Research Grant (CRG/2021/002150) from SERB, Department of Science and Technology (DST), Government of India, and IISER-K intramural funding to AG. The Pre-doctoral fellowship for Ruturaj is supported by Intramural Institute funding (IISER-K). SM is supported by a pre-doctoral fellowship from Council of Scientific and Industrial Research (CSIR), India. We thank Svetlana Lutsenko (Johns Hopkins University) for sharing the HA-ATP7A construct and Carolyn Machamer (Johns Hopkins University) for sharing antibodies with us.

## Author contributions

AG, Ruturaj, ERB and RS designed the experiments. AG, ERB and Ruturaj wrote the manuscript. Ruturaj, MM, RS, SS and SM did the experiments and analyzed the data. SM wrote the codes and analyzed the data. All authors reviewed the results and approved the final version of the manuscript.

The authors declare no competing financial interests.

**Video1. Representative video of dispersion rate of ATP7A from Golgi in response to copper**. MDCK cells transfected with mKO2-HA-ATP7A were plated on glass bottom confocal dishes. Images of sub-confluent MDCK cells treated with 50µM CuCl_2_ were captured at an interval of 60 seconds for 1 hour. Pixels having 95% of the signals under basal conditions were considered as Golgi, the disappearance of which represents the Golgi-exit rate.

**Video2. Representative video of dispersion rate of ATP7B from Golgi in response to copper**. MDCK cells transfected with eGFP-ATP7B were plated on glass bottom confocal dishes. Images of sub-confluent MDCK cells treated with 50µM CuCl_2_ were captured at an interval of 60 seconds for 1 hour. Pixels having 95% of the signals under basal conditions were considered as Golgi, the disappearance of which represents the Golgi-exit rate.

**Video3. Representative video of dispersion rate of ATP7B ΔF37-E45 mutant from Golgi in response to copper.**

MDCK cells transfected with eGFP-ATP7B ΔF^37^-E^45^ were plated on glass bottom confocal dishes. Images of sub-confluent MDCK cells treated with 50µM CuCl_2_ were captured at an interval of 60 seconds for 1 hour. Pixels having 95% of the signals under basal conditions were considered as Golgi, disappearance of which represents the Golgi-exit rate.

---

**Figure S1.**
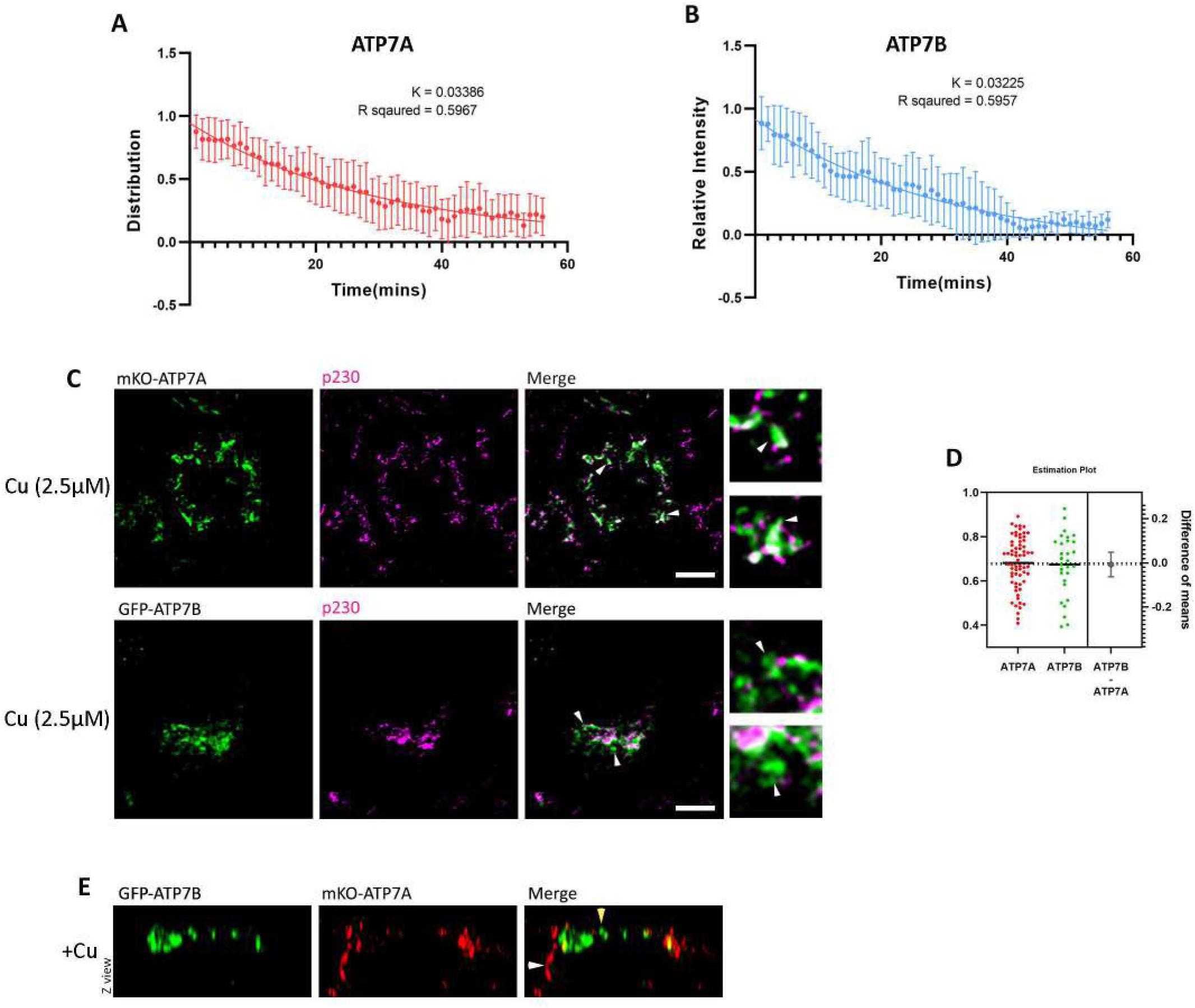
**(A)** Fitted curve of decreased pixel count of mKO2-HA-ATP7A showing its exit rate in response to copper (i.e., k value = 0.03386). **(B)** Fitted curve of decreased pixel count of GFP-ATP7B showing its exit rate in response to copper (i.e., k value = 0.03225). **(C)** Representative confocal images of CuCl_2_ (10µM; 30mins) treated cells transfected with either mKO2-HA-ATP7A or GFP-ATP7B and co-stained with p230. (Arrowheads indicate TGN exited proteins) **(D)** Estimation plot of Fig. 1E created in GraphPad Prism v9.4. **(E)** Confocal images showing co-transfected mKO2-HA-ATP7A and GFP-ATP7B under copper-treated conditions. Yellow arrow denotes ATP7B localizing at the apical surface. White arrows mark ATP7A at the basolateral surface. Scale bar, 5µm.

**Figure S2.**
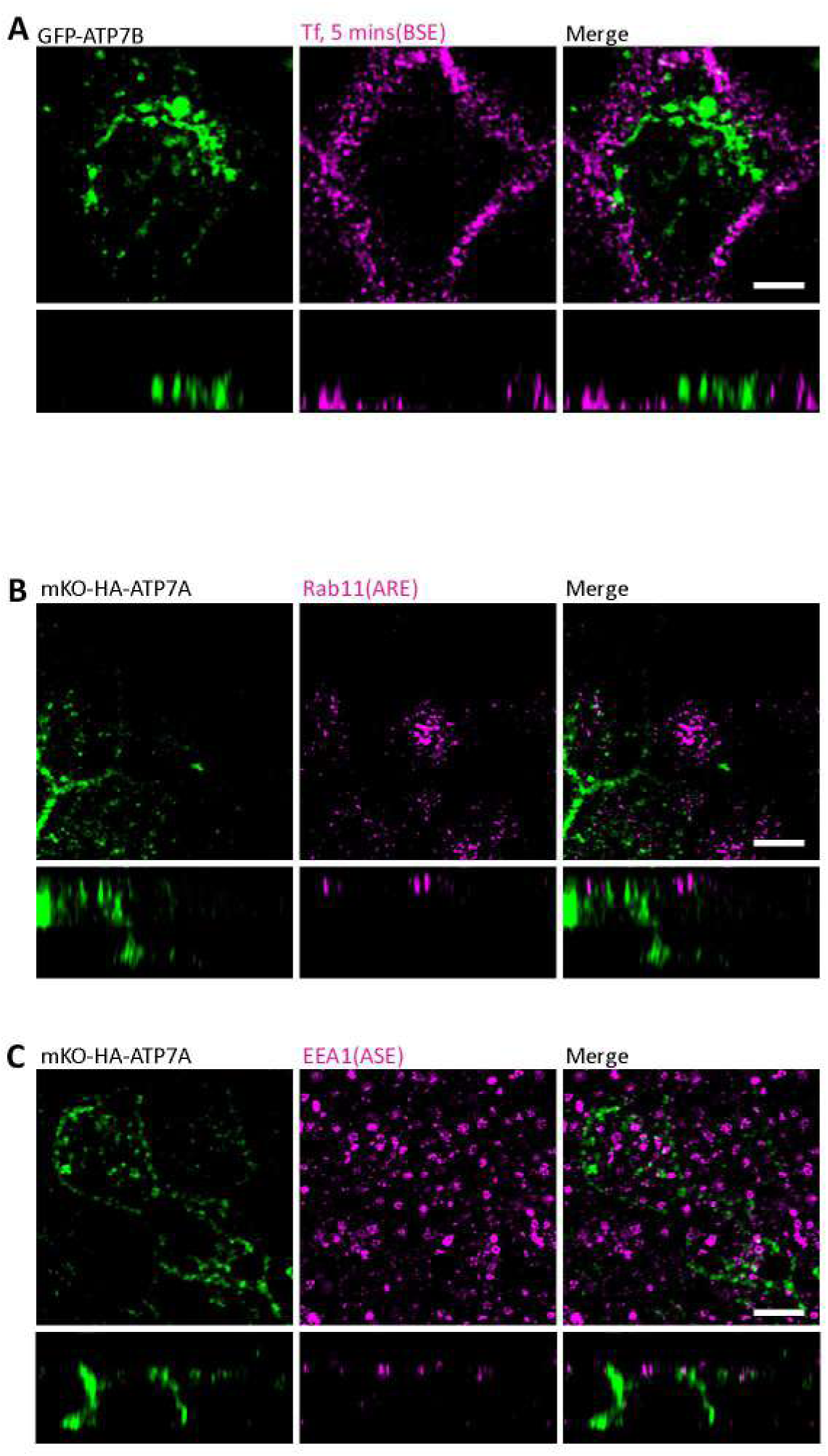
**(A)** Confocal image of GFP-ATP7B (green) co-stained for BSE (magenta; 5mins Tf Internalization) shows no localization of ATP7B in BSE. **(B)** Confocal image of mKO-HA-ATP7A (green) co-stained with Rab11 (magenta) shows no localization of ATP7A in ARE (Rab11 positive compartment). **(C)** Confocal image of mKO-HA-ATP7A co-stained with EEA1 shows no localization of ATP7A in ASE (EEA1 positive compartment). Scale bar, 5µm.

**Figure S3.**
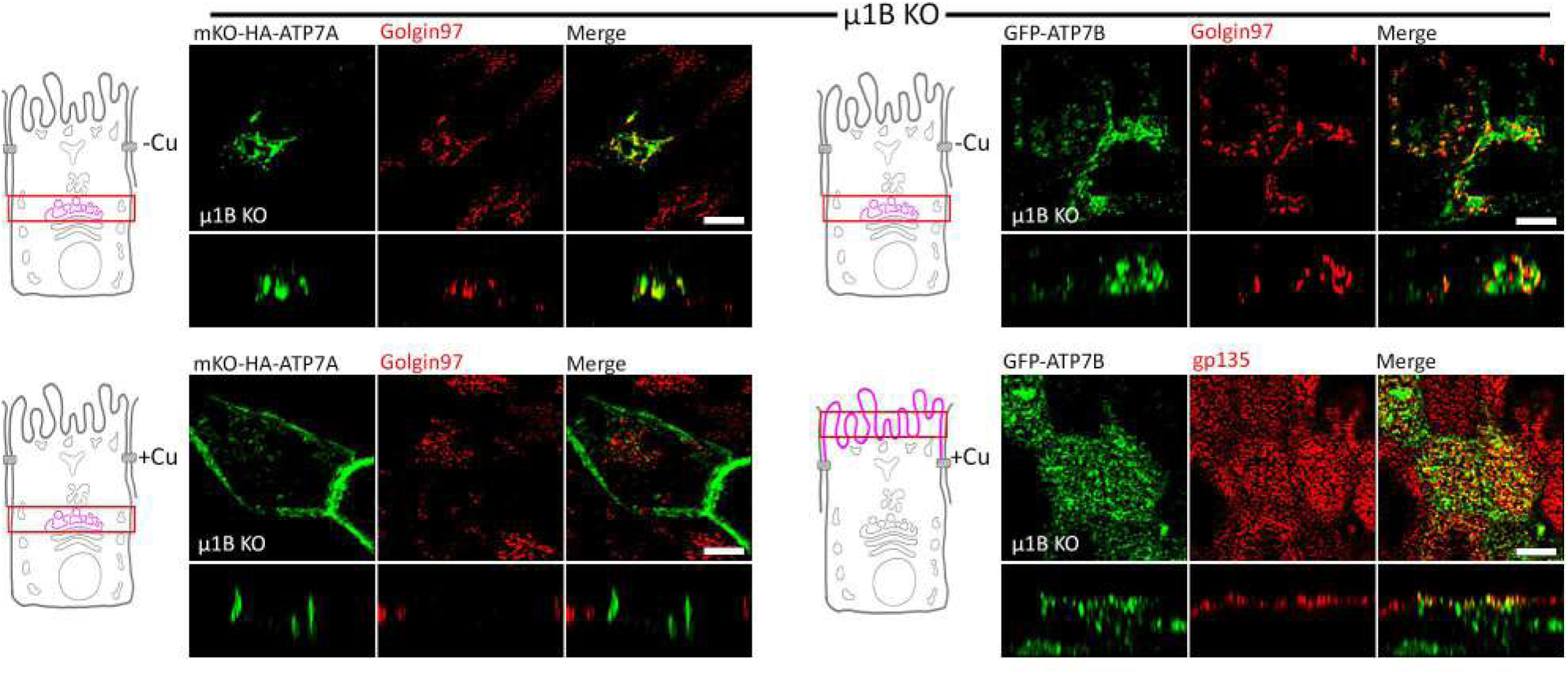
Confocal image of cells either transfected with mKO-HA-ATP7A or GFP-ATP7B in AP1B KO cells (i.e., µ1B KO cells). Under copper deprived as well as copper-treated conditions both the ATPases exhibit wild-type-like phenomena. Scale bar, 5µm.

**Figure S4.**
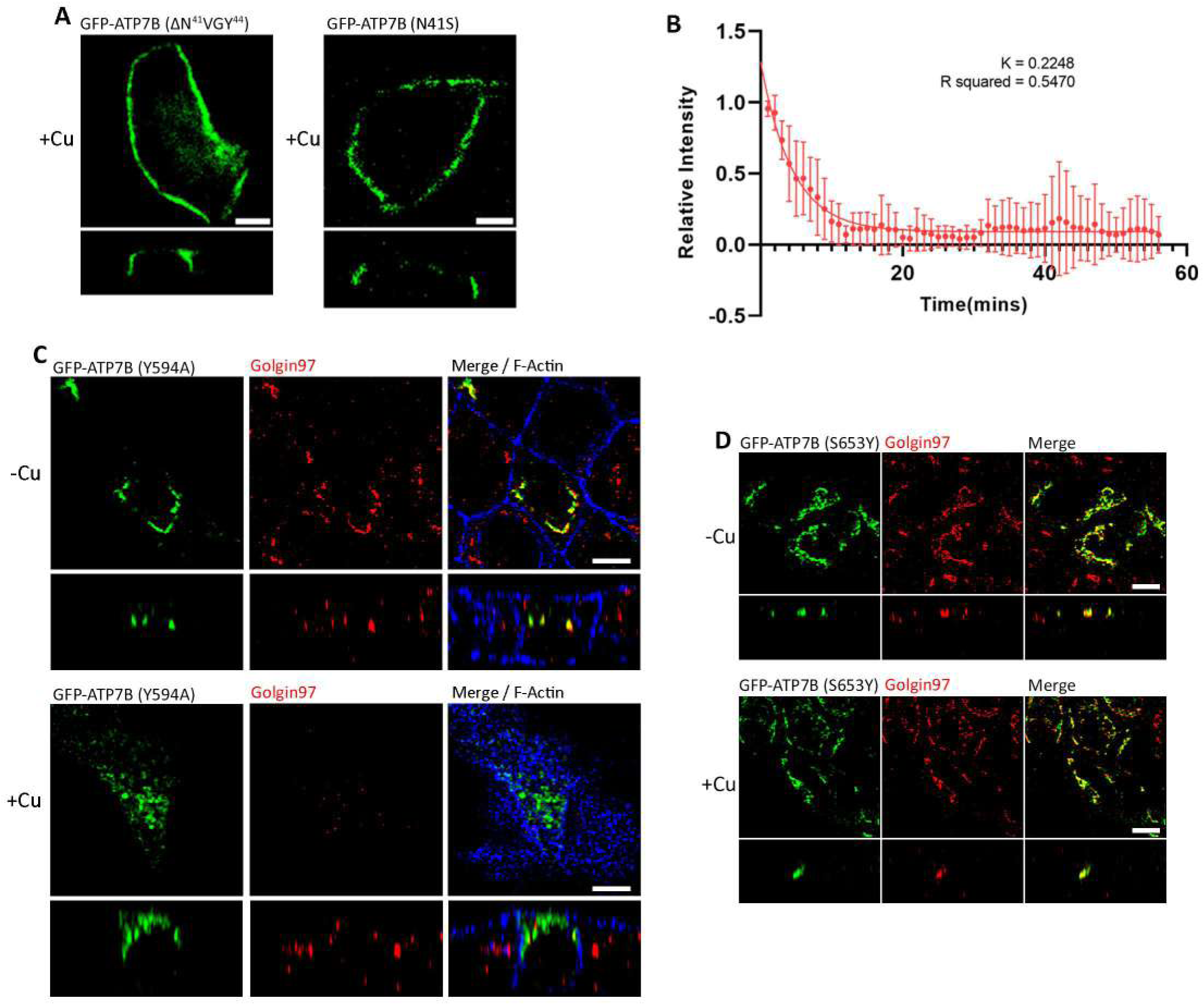
**(A)** Confocal images of ΔN^41^VGY^44^ and N41S of ATP7B mutants shows their basolateral localization in copper-treated conditions. **(B)** Fitted curve of decreased pixel count of Δ9aa GFP-ATP7B mutant in response to copper showing its dispersion and the exit rate (i.e., k value = 0.2248). **(C)** Confocal images of Y594A-ATP7B mutant shows wildtype-like copper-mediated trafficking behaviour. **(D)** Confocal image of S653Y-ATP7B mutant unable to exit TGN in copper-treated condition. Scale bar, 5µm.

